# Native holdup (nHU) to measure binding affinities from cell extracts

**DOI:** 10.1101/2022.06.22.497144

**Authors:** Boglarka Zambo, Bastien Morlet, Luc Negroni, Gilles Trave, Gergo Gogl

## Abstract

Characterizing macromolecular interactions is essential for understanding cellular processes, yet nearly all methods used to detect protein interactions from cells are qualitative. Here, we introduce the native holdup (nHU) approach to quantify equilibrium binding constants and explore binding mechanisms of protein interactions from cell extracts. Compared to other pulldown-based assays, nHU requires less sample preparation and can be coupled to any analytical methods, such as western blotting (nHU-WB) or mass spectrometry (nHU-MS) as readouts. We use nHU to explore interactions of SNX27, a cargo adaptor of the retromer complex and find good agreement between *in vitro* affinities and those measured directly from cell extracts using nHU. This challenges the unwritten paradigm stating that biophysical parameters like binding constants cannot be accurately determined from cells or cellular extracts. We discuss the strengths and limitations of nHU and provide simple protocols that can be implemented in most laboratories.

## Intro

Quantitative high-throughput (HTP) approaches are needed to explore the *human protein-protein affinity interactome* ^1 2 3^. Commonly, protein complexes are identified from cellular extracts with pulldown-based approaches involving a bait immobilized on beads via antibodies (immunoprecipitation) or affinity tags (affinity purification). Protein complexes are then purified from cell extracts on the beads by applying extensive washing protocols that reduce non-specific background step-wise while retaining the bound specific interaction partners ^4 5^. Other HTP methods used to discover protein complexes are based on two-hybrid screens, the co-fractionation of molecular assemblies, display technologies, or the irreversible modification of proteins found in close proximity, and are often coupled to a pulldown-based approach ^6 7 8 9 10^. While all of these methods are powerful for discovering interactions, they are qualitative and do not shed light on the biophysical attributes underpinning the observed interactions, such as dissociation constants (*K*_d_) or its negative logarithm (p*K*_d_).

Molecules neither “interact” nor “not interact”, rather their interactions follow physical rules. For example, the law of mass-action in binding equilibrium defines the degree of complex formation as a function of binding affinities and cellular concentrations. Quantitative assays aiming to measure binding affinities at HTP are mostly limited to fragmentomic approaches, where interactions are studied between minimal binding fragments, most usually between globular domains and peptide motifs. Most recently, we characterized several thousands of domain-motif affinities using the holdup assay, a single-point binding experiment that measures the degree of complex formation under equilibrium ^11 12^. These and other recent advances brought proteome-wide fragmentomic affinity screening of elementary reactions within reach ^13 14 15 16^. However, fragmentomic approaches do not reveal how affinities change when minimal binding fragments are embedded in the full-length proteins or even larger macromolecular complexes.

As a solution, we developed the quantitative native holdup (nHU) assay to measure binding affinities of full-length proteins directly from native cell extracts. We demonstrate that nHU can be coupled to various protein analytical methods, such as western blot or mass spectrometry, exploiting all advances of protein analytics. We explore the interactions of Sorting Nexin 27 (SNX27), a component of the retromer complex involved in the endosome-to-plasma membrane protein recycling ^17 18^. We find that the nHU assay is robust and reliable in quantifying equilibrium binding constants over a wide affinity range. We show that nHU and fragmentomics are highly complementary as affinities measured with these approaches are related but not necessarily identical, due to the presence of many other factors in full-length proteins, such as multivalency or conformational heterogeneity.

## Results

### Principles of nHU

In a holdup experiment, an analyte solution is incubated with either control or bait-saturated resin and the differential depletion of prey protein within the liquid phase fraction is measured with analytical methods to calculate the equilibrium binding constant ^19 11 12^. We used streptavidin resin saturated with biotinylated peptides or proteins as baits and biotin or biotinylated-MBP as controls. The saturated resin is equilibrated with the analyte and the liquid phase is rapidly separated from the resin by either fast filtration or simply by pipetting after a short centrifugation. The unbound prey fraction is quantified by the relative prey concentration of the bait and control samples and the complementary bound fraction, conventionally called binding intensity (BI), is determined as follows:

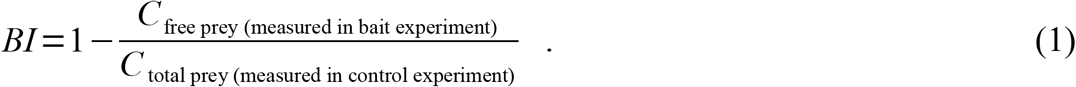

In nHU, total cell extracts are used as an analyte and BI values of preys are measured with quantitative protein analytical methods, such as western blot or mass spectrometry (Figure 1A) ^19 20^. The qualitative proof-of-concept of nHU was demonstrated before and here we develop it into a quantitative assay for the first time ^19^. Cell extracts contains thousands of distinct preys, all present at very low concentrations. We assume that even the cumulative amount of bound prey fractions occupies only a minor, negligible fraction of the total amount of the immobilized bait. Taken together, the apparent equilibrium constants for interactions following a simple bimolecular binding mechanism can be calculated using the hyperbolic binding equation:

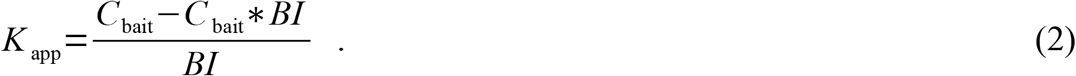

**Figure 1.**
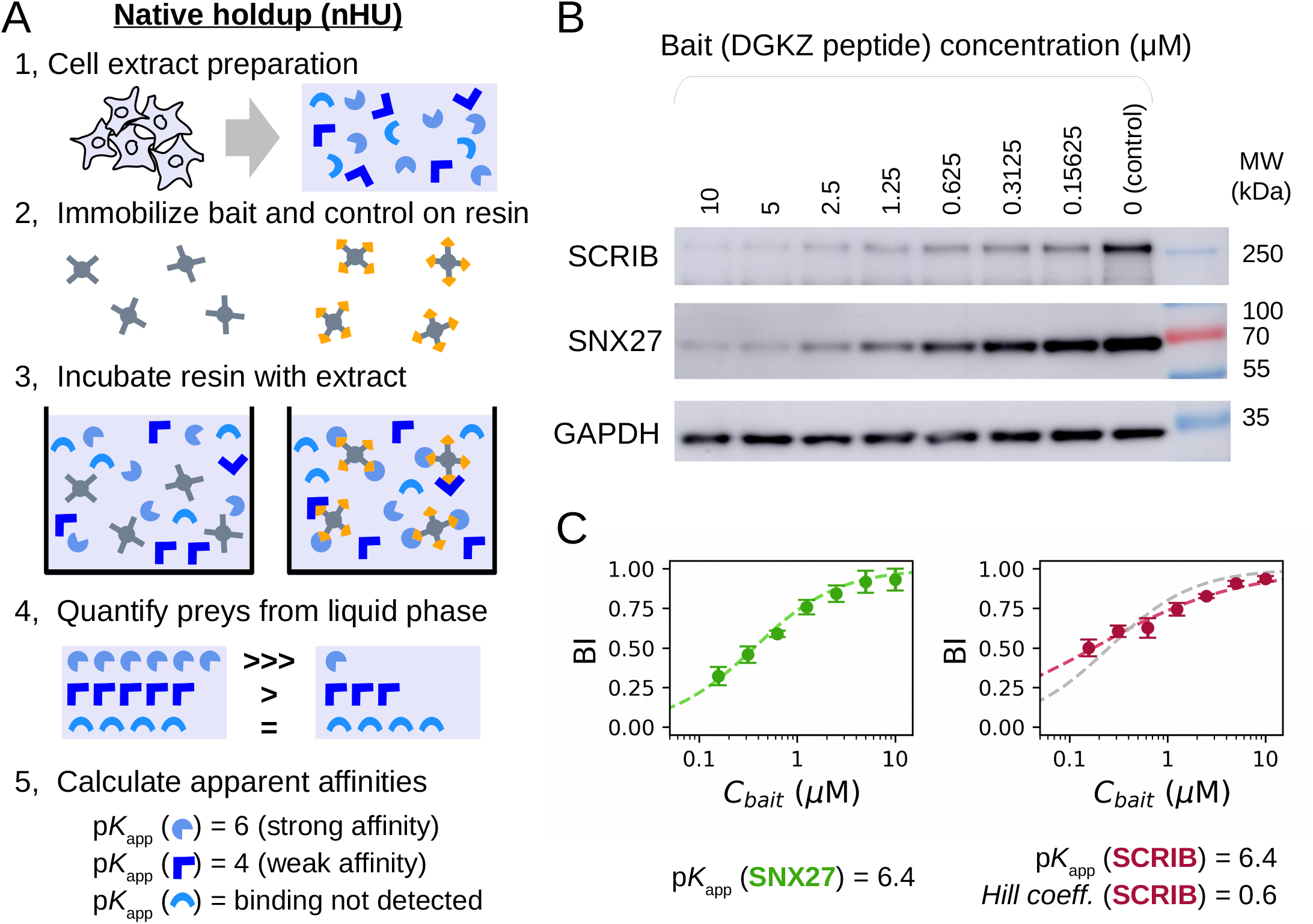
Principle and simple demonstration of nHU. (A) Schematic pipeline of nHU. Biotinylated baits and controls are immobilized on streptavidin resin at high concentration and are mixed with cellular extracts. After the binding equilibrium is reached, the liquid phase is separated by filtration or by centrifugation and amounts of prey proteins are determined using standard protein analytical tools, such as western blot or mass spectrometry. The measured concentration ratio, in combination with the estimated amount of the immobilized bait concentration can be directly converted into apparent equilibrium dissociation constants (p*K*_app_). Note that the discarded resin from step 3 can be optionally processed as a regular pulldown experiment. (B) Demonstration of nHU titration experiment using the biotinylated PBM peptide of DGKZ as bait and biotin as control. Increasing amounts of bait-saturated resins were incubated with total Jurkat extracts. Supernatant fractions were probed with specific antibodies against endogenous full-length PDZ-domain-containing proteins SCRIB and SNX27. (C) Results of nHU-WB experiments presented in panel B. BI values of SNX27 (green, left) and SCRIB (red, right) were first fitted with a hyperbolic binding equation (dashed line). In the case of SCRIB, significantly better fit was obtained using the Hill equation (red dashed line) compared to a hyperbolic binding equation (gray dashed line). Determined parameters are indicated below the plots. BI values were determined based on 3 replicates. See Supplementary Figure 1 and Supplementary Table 1 for additional data.

The accuracy of the calculated affinities depends on the accuracy of the estimation of the concentration of the immobilized bait (*C*_bait_). This concentration needs to be higher than the cumulative concentration of all prey molecules of the extract that are prone to be captured by the bait. Previously, we determined the binding capacity of streptavidin resin for various ligands by substituting affinities measured with orthogonal methods in binding equations such as Eq. 2. As a rule of thumb, if 50 μl saturated streptavidin resin (Streptavidin Sepharose High Performance, Cytiva) is mixed with 200 μl extract, the estimated *C*_bait_ is around 10 μM ^12 21^.

### Measuring binding affinity with nHU coupled to western blot (nHU-WB)

To demonstrate that nHU can be used to measure affinities of a full-length protein directly from a cell extract, we chose to study an already described interaction between the PDZ-binding motif (PBM) peptide of diacylglycerol kinase zeta (DGKZ) and full-length SNX27 endogenously present in Jurkat extract ^22^. First, we saturated streptavidin resin with biotin or biotinylated peptide. Then, we incubated total Jurkat cell extracts with various mixes of control and bait saturated resin by always keeping the resin/analyte ratio constant. This way, the concentration of SNX27 is fixed (determined by the lysate) and the concentration of the immobilized DGKZ peptide covers a wide range of concentration ^23 24^. The supernatant fractions of each experiment were assayed by western blot using a specific antibody against SNX27 (Figure 1B, Supplementary Figure 1A). As expected, the measured BI values of SNX27 decreased when *C*_bait_ was decreased, following the hyperbolic binding equation (Eq. 2) revealing an affinity of 6.4 p*K*_d_ (Figure 1C, left panel; Supplementary Table 1). This interaction between the PBM peptide of DGKZ and the isolated PDZ domain of SNX27 was already studied by both calorimetry and fragmentomic holdup and the affinities were found to be 5.7 and 6.0 p*K*_d_, respectively ^12 22^. Thus, we found nHU to be a robust and versatile method for determining biophysical properties of an endogenous full-length protein directly from cellular extract.

### Single-point nHU-WB to measure motif-protein affinities in medium-throughput

Titration experiments are key for investigating precise binding mechanism. However, they come with low throughput and high experimental cost. For this reason, we used single-point holdup experiments to measure affinities of thousands of fragmentomic interactions at HTP in our previous study ^12^. Similarly, we propose that nHU experiments can also be performed using a single bait concentration to probe binding equilibrium by assuming the binding mechanism. We used 12 different biotinylated 10-mer PBM peptides as baits or biotin as a negative control and performed nHU at a single bait concentration (10 μM) using total Jurkat extracts as analyte (Figure 2A, Supplementary Figure 1B). We measured the depletion of SNX27 from the supernatants using nHU-WB and the measured binding intensities of the PBM-baits were converted into affinities using Eq. (2). The resulting affinities between full-length SNX27 and the 12 peptide baits showed excellent agreement with their fragmentomic affinities measured with the isolated PDZ domain of SNX27 with a Pearson correlation coefficient (PCC) of 0.95 (Figure 2B, left panel; Supplementary Table 1) ^12^.

**Figure 2.**
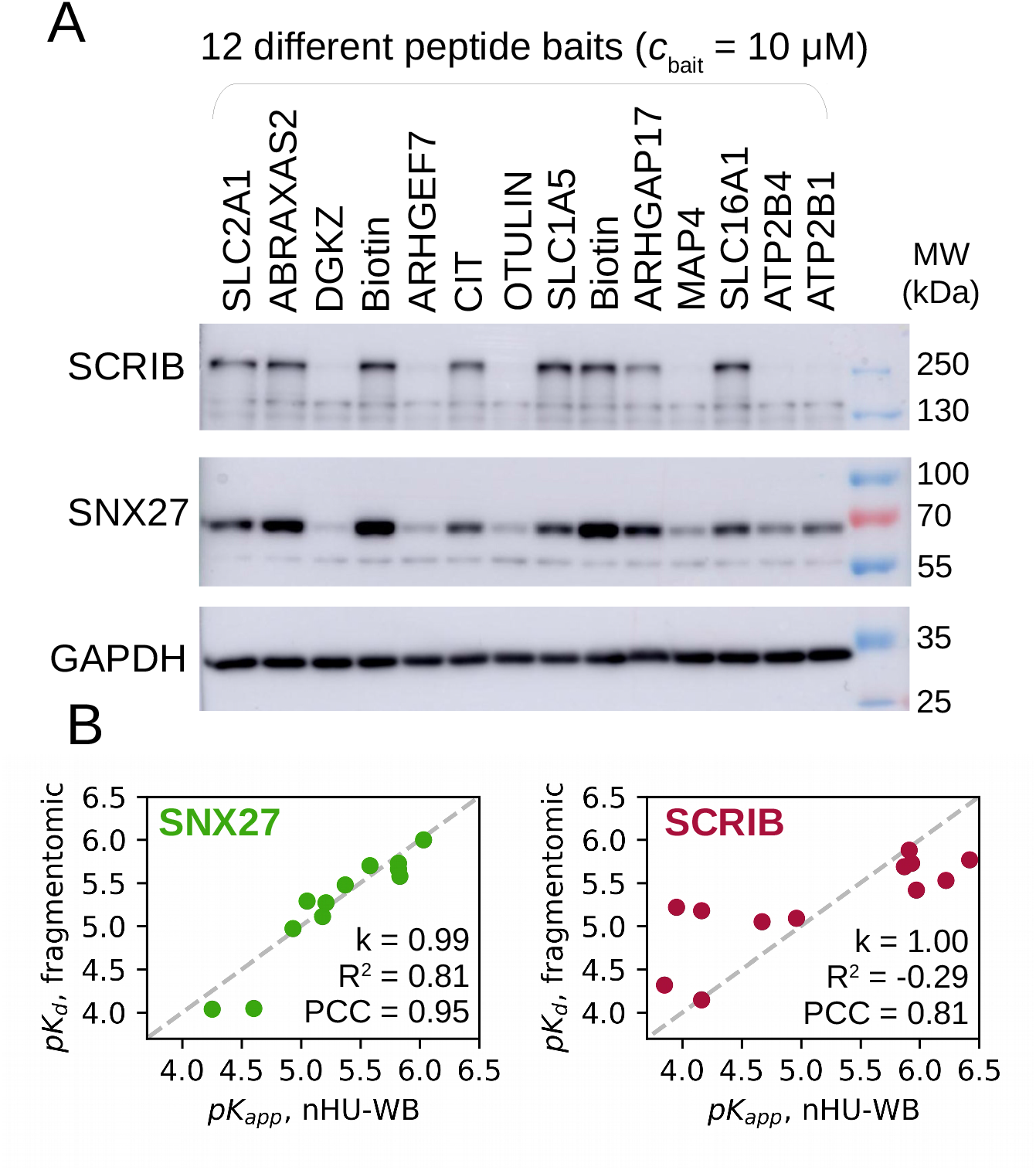
Single-point nHU for rapid apparent affinity measurements. (A) Demonstration of single-point nHU-WB using 12 different biotinylated PBM peptides (baits) or biotin (control) saturated streptavidin resin and total Jurkat extracts. Supernatant fractions were probed with specific antibodies against endogenous full-length PDZ-domain-containing proteins SCRIB and SNX27. (B) Results of the nHU-WB experiment presented in panel A. Correlation between *in vitro* fragmentomic affinities measured using PBM peptides and isolated PDZ domains ^12^ and apparent affinities measured with nHU between PBM peptides and full-length proteins. In the case of SCRIB (on right), the site-specific fragmentomic affinities were combined assumed additivity. Direct proportionality was assumed between affinities (gray dashed line) and the coefficient of proportionality (k), R^2^ values and Pearson correlation coefficient (PCC) values are indicated. Affinities were determined based on 3 replicates. See Supplementary Figure 1 and Supplementary Table 1 for additional data.

However, not all interactions follow simple binding mechanism ^25^. For example, the 4 PDZ domains of Scribble (SCRIB) can synergize and its interactions with PBMs are neither bimolecular nor single-sited. In addition to SNX27, we also measured the depletion of endogenous full-length SCRIB in the previous nHU-WB experiments. In the titration experiment, the affinity between the DGKZ peptide and full-length endogenous SCRIB was found to be more than an order of magnitude stronger than any of the site-specific affinities of its isolated domains (6.4 p*K*_d_ vs. 5.1, 4.9, 4.7, 4.4 p*K*_d_) (Figure 1C, right panel; Supplementary Table 1) ^26^. We also found that the interaction displays negative cooperativity with a Hill coefficient of 0.6 ^27^. Consequently, neither single-point holdup experiments, nor Eq. 2. can not reveal site-specific affinities of SCRIB with absolute confidence. Still, they could be used to calculate apparent affinities for ranking different interaction partners. Site-specific fragmentomic affinities can be combined to approximate an additive affinity of all interaction sites ^12^. Despite these rough approximations, a good correlation (PCC = 0.81) was found between the apparent affinities measured by single-point nHU and the combined fragmental affinities (Figure 2B, right panel; Supplementary Table 1). Therefore, single-point nHU experiments were found to be robust to explore intrinsic properties of a large number of interactions, regardless of their binding mechanisms.

### nHU coupled to mass spectrometry (nHU-MS) to measure domain-protein affinities proteome-wide

We coupled nHU to label-free quantitative mass spectrometry in order to quantify affinities of the SNX27 PDZ domain (SNX27_PDZ) proteome-wide (Figure 3A, Supplementary Table 1). We used this isolated PDZ domain fused to a biotinylated-His_6_-MBP-tag as the bait or the tag alone as a control. To show the enrichment of preys on the resin that was depleted from the supernatant, the resin was further processed after the nHU reaction according to a standard pulldown protocol (Figure 3B).

**Figure 3.**
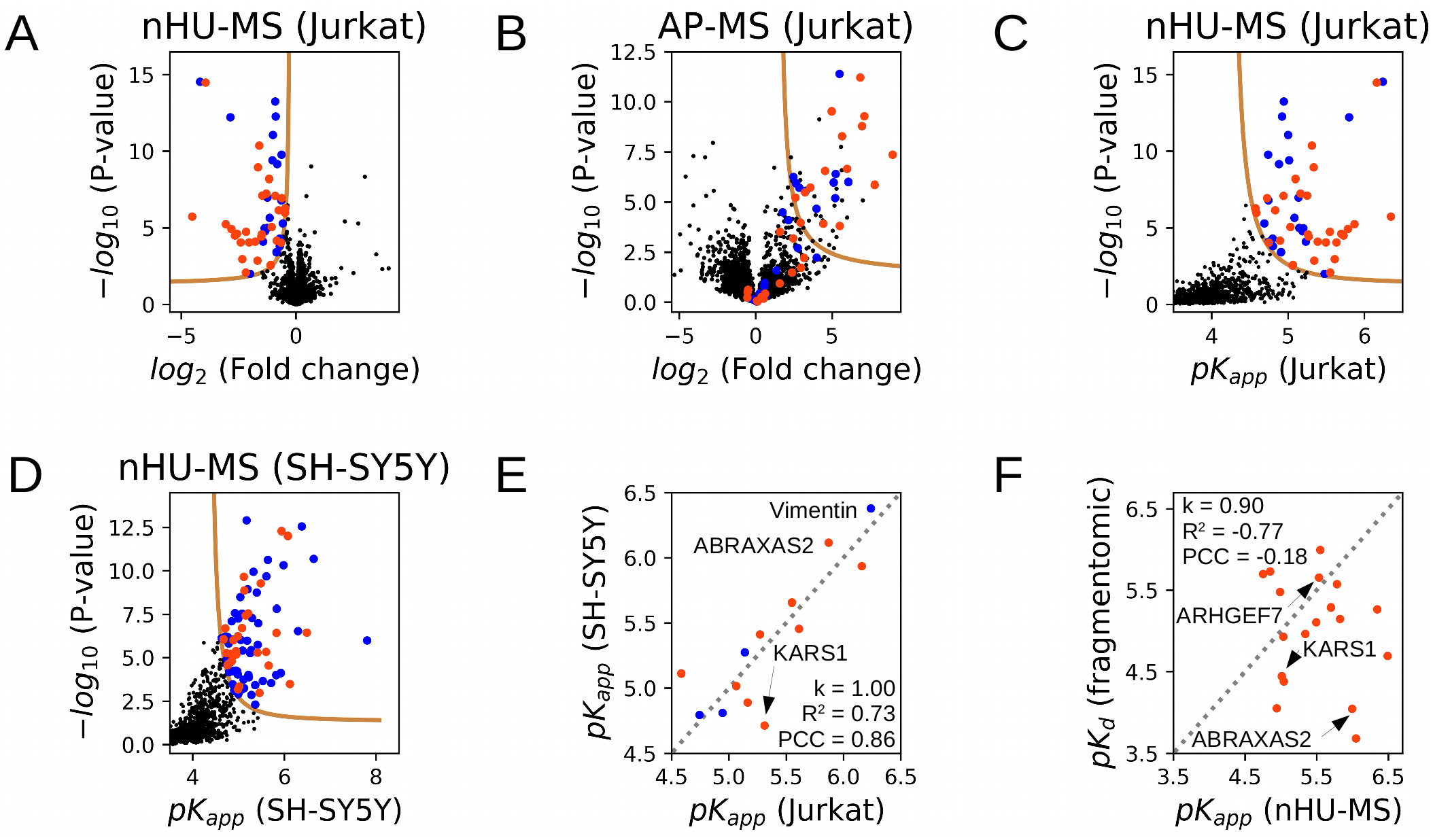
nHU-MS to survey affinities of the SNX27 PDZ domain proteome-wide. (A) Volcano plot of the nHU-MS experiment performed with SNX27_PDZ bait on total Jurkat extracts (n=6). Identified interaction partners with or without C-terminal putative PBMs are colored with orange or blue, respectively. (B) Volcano plot of the control pulldown experiment performed on the leftover resin of the nHU experiment (n=6). Preys are colored according to their coloring on panel A. (C and D) Converted affinities of nHU experiments measured with Jurkat (C) or SH-SY5Y (D) extracts. (A-D) P-values were calculated using a two-sided unpaired T-test for nHU-MS and AP-MS experiments. The statistical significance thresholds for binding (tan lines) were determined at 1 σ and > 0.05 P-value. (E) Correlation between apparent affinities of SNX27_PDZ measured with Jurkat or SH-SY5Y extracts. (F) Correlation between *in vitro* fragmentomic affinities measured using PBM peptides and the isolated PDZ domain of SNX27 ^12^ and apparent affinities measured with nHU between the same domain and full-length proteins (average affinities measured from the two cell extracts). Direct proportionality was assumed between affinities and the coefficient of proportionality (k), R^2^ values and Pearson correlation coefficient (PCC) values are indicated on panels E and F. Gray dashed line indicates the diagonal on panel E and F. See Supplementary Figure 2 and Supplementary Table 1 for additional data.

We performed the experiment using Jurkat and SH-SY5Y cell extracts (Figure 3C, D, respectively; Supplementary Figure 2A). In total, the affinities were assayed against 3,548 full-length endogenous proteins, out of which 121 showed statistically significant depletion, most of which were also enriched on the resins during pulldown. The determined apparent affinities for the 13 interaction partners of SNX27_PDZ that we quantified from both cell extracts were directly proportional with a PCC of 0.86 (Figure 3E). We identified 52 proteins with putative C-terminal PBMs among the interaction partners of SNX27_PDZ. In our previous fragmentomic screen, the binding affinities of >400 isolated PBM peptides were already assayed with SNX27_PDZ ^12^. From the 52 full-length interaction partners, the PBMs of 19 were included in this array and with one exception, all of these interactions showed quantifiable *in vitro* affinity. Apart from this qualitative comparison, these fragmentomic affinities showed no correlation with the apparent affinities of nHU (PCC = -0.18) (Figure 3F). While some partners, like ARHGEF7, have very similar affinities when measured using their C-terminal fragments, or when measured from cell extracts, most interactions were stronger in nHU experiments. On average, affinities were found to be >3-fold stronger (Δp*K* ≈ 0.5) when measured using full-length proteins in nHU compared to the affinities of isolated PBM peptides.

### nHU-MS to measure protein-protein affinities proteome-wide

We performed a nHU-MS experiment using total Jurkat extracts as analyte and recombinant full-length SNX27 fused to a biotinylated-His_6_-MBP-tag as a bait or the tag alone as a control (Figure 4A, B, Supplementary Table 1). In addition to the PDZ domain, full-length SNX27 contains a Phox homology (PX) and a FERM (4.1, ezrin, radixin, moesin) domain. We quantified 3,128 full-length endogenous preys, out of which 198 showed statistically significant depletion in nHU. Most of these targets were also enriched in a subsequent control pulldown experiment (Supplementary Figure 2B). Among the interaction partners of SNX27, 26 proteins were also identified in the nHU experiments as partners of SNX27_PDZ, including 13 with putative C-terminal PBMs, and the obtained affinities correlate with a PCC of 0.66 (Figure 4C).

**Figure 4.**
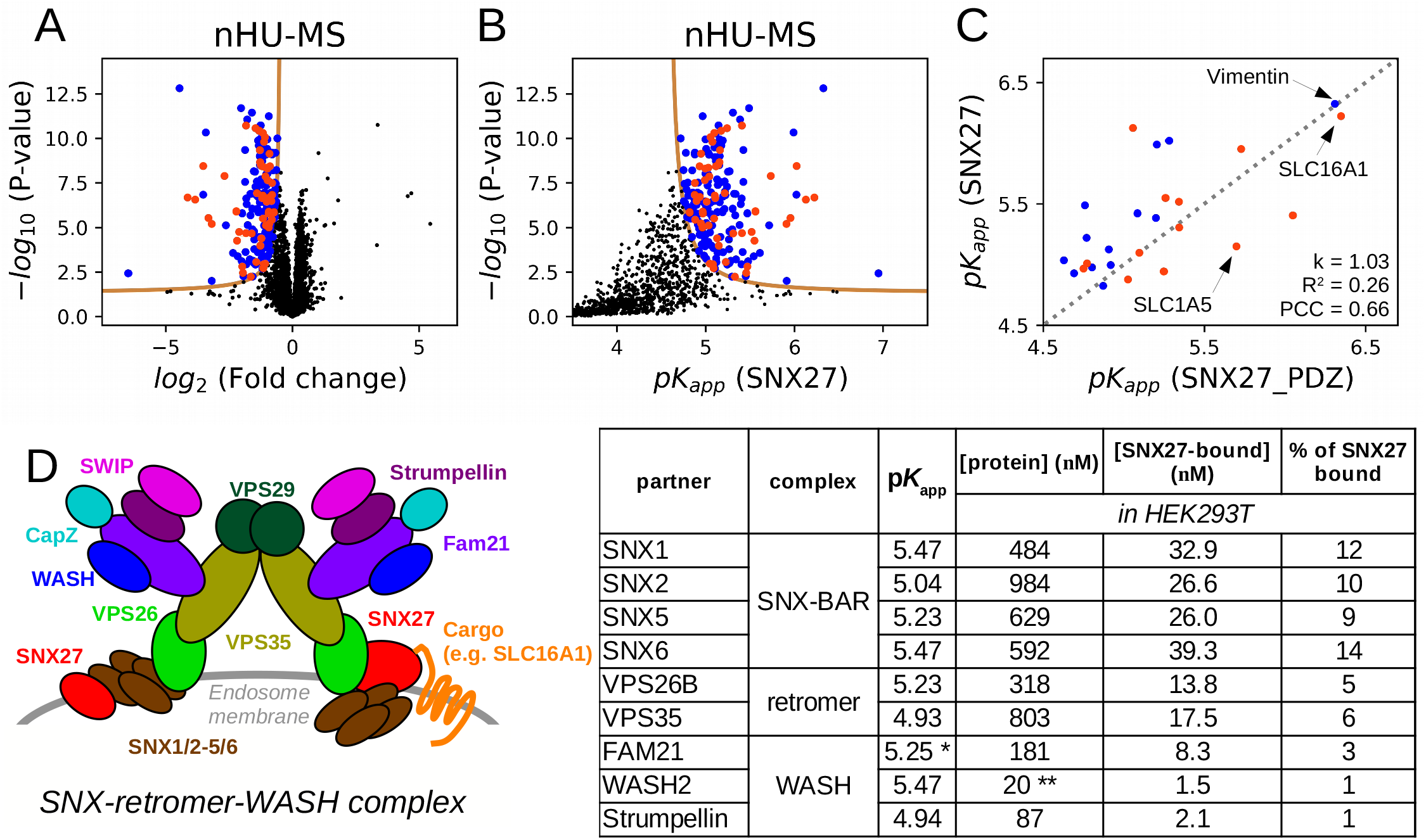
nHU-MS to survey affinities of full length SNX27 proteome-wide. (A, B) nHU-MS experiment performed with recombinant SNX27 bait and total Jurkat extracts (n=6) analyzed as a function of fold change (A) or converted affinities (B). Identified interaction partners with or without C-terminal putative PBMs are colored with orange or blue, respectively. P values were calculated using a two-sided unpaired T-test and statistical thresholds for binding (tan lines) were determined at 1 σ and > 0.05 P. (C) Correlation between apparent affinities of the SNX27_PDZ or full length SNX27. Direct proportionality was assumed between affinities and obtained parameters as well as the diagonal (gray dashed line) are indicated. (D) Coarse topology, measured affinities and estimated steady state of the SNX-retromer-WASH complex. The two hypothetical topological position of SNX27 in the complex based on current observations were shown on the opposite sides of the dimeric complex. Measured affinities of SNX27 were combined with estimated total protein concentrations of HEK293T cells (taken from OpenCell^35^) using the quadratic binding equation to estimate the amount of SNX27-bound complexes. Note that amounts of complexes of the same sub-complexes are in the same regime: ∼10-15% of the total amount of SNX27 found in HEK293T cells (280 nM) is bound to SNX-BARs, ∼5% of SNX27 is bound to retromer and ∼2% of SNX27 is bound to WASH. *Affinity determined for FAM21 showed statistical significance below threshold. ** The concentration of WASH2 in HEK293T is unknown and is substituted with the concentration of WASH1. See Supplementary Figure 2 and Supplementary Table 1 for additional data.

Affinities obtained from nHU can originate either from direct or indirect interactions through large complexes (Supplementary Figure 2C). We identified the retromer complex as an interaction partner of SNX27 with the strongest affinity measured with VPS26B ^28 29^. We also detected the association of SNX27 and the heterodimeric SNX1/2-SNX5/6 SNX-BAR complex with the strongest affinity measured with SNX1. A short fragment of SNX1/2 was reported to interact with the FERM domain of SNX27 with comparable affinity to the one obtained from nHU ^30^. We identified the actin-regulatory Wiskott Aldrich Syndrome protein and scar homologue (WASH) complex as the partner of SNX27 with the strongest affinities measured with WASH2 and FAM21 ^31^. The 1.5 mDa multi-tRNA synthetase complex co-fractionates with the retromer complex in extracts, and we found that all components have detectable affinity with SNX27_PDZ with the strongest interaction with KARS1 ^32^. We identified the 320 kDa BRCC36 isopeptidase (BRISC) complex as a target of SNX27_PDZ with the strongest affinity measured with ABRAXAS2 ^33^. Finally, we also detected the oligomeric small GTPase-regulator GIT-PIX complex as the interaction partner of SNX27_PDZ with ARHGEF7 (also called β-PIX) displaying the strongest affinity ^34^. In the case of the last three complexes, that were identified as the partner of SNX27_PDZ, only the partners with highest affinities had PBMs satisfying the SNX27_PDZ consensus. In these examples, it appears that when large complexes are captured by a bait in nHU assays the subunit that binds the bait directly displays the strongest measured affinity, while weaker affinities of other subunits can be an asset to map topologies of the complexes.

Affinities obtained from nHU experiments can be used to estimate steady states of networks in various conditions by combining with cellular concentrations of proteins. We took apparent affinities between SNX27 and other components of the SNX-retromer-WASH complex and combined them with estimated total protein concentrations measured for HEK293T cells using the quadratic binding equation (Figure 4D) ^35^. Based on this coarse analysis, 10% of cellular SNX27 is expected to bind to SNX-BARs, 5% to the retromer complex and 2.5% to the WASH complex in HEK293T cells.

### Interactions of SNX27 with membrane and filamentous proteins

For further mechanistic characterization, we selected three representative full-length interaction partners of SNX27 that displayed similar affinities with full-length SNX27 or SNX27_PDZ baits and were thus likely to be specific interaction partners of the PDZ domain (Supplementary Figure 4). The monocarboxylate transporter 1 (SLC16A1) ^36^, and the intermediate filament-forming Vimentin ^37^ were among the strongest partners of SNX27 and the neutral amino acid transporter B(0) (SLC1A5) ^38^ displayed weaker affinity. To explore the binding mechanisms of these interactions, we performed a nHU-WB titration experiment with SNX27 (Figure 5, Supplementary Figure 3, Supplementary Table 1). We have found that SLC1A5 followed a simple binding mechanism (as in Eq. 2). Although SLC16A1 also followed similar binding mechanism, approximately 10% of the detected protein seemed to be incapable to binding, possibly due to regulatory modifications of its C-terminal tail segment. In the case of filamentous Vimentin, we observed a positive cooperative binding mechanism resulting in a stronger SNX27 binding affinity than that approximated from single-point measurements ^27^. However, in spite of the more complex binding mechanisms of Vimentin and SLC16A1, their affinities estimated from single point nHU experiments were still indicative for their strong interaction with SNX27. It is worth mentioning that characterizing interactions of full-length transmembrane or filamentous proteins with traditional quantitative biochemical methods is highly challenging, but seems to be easily addressed with nHU.

**Figure 5.**
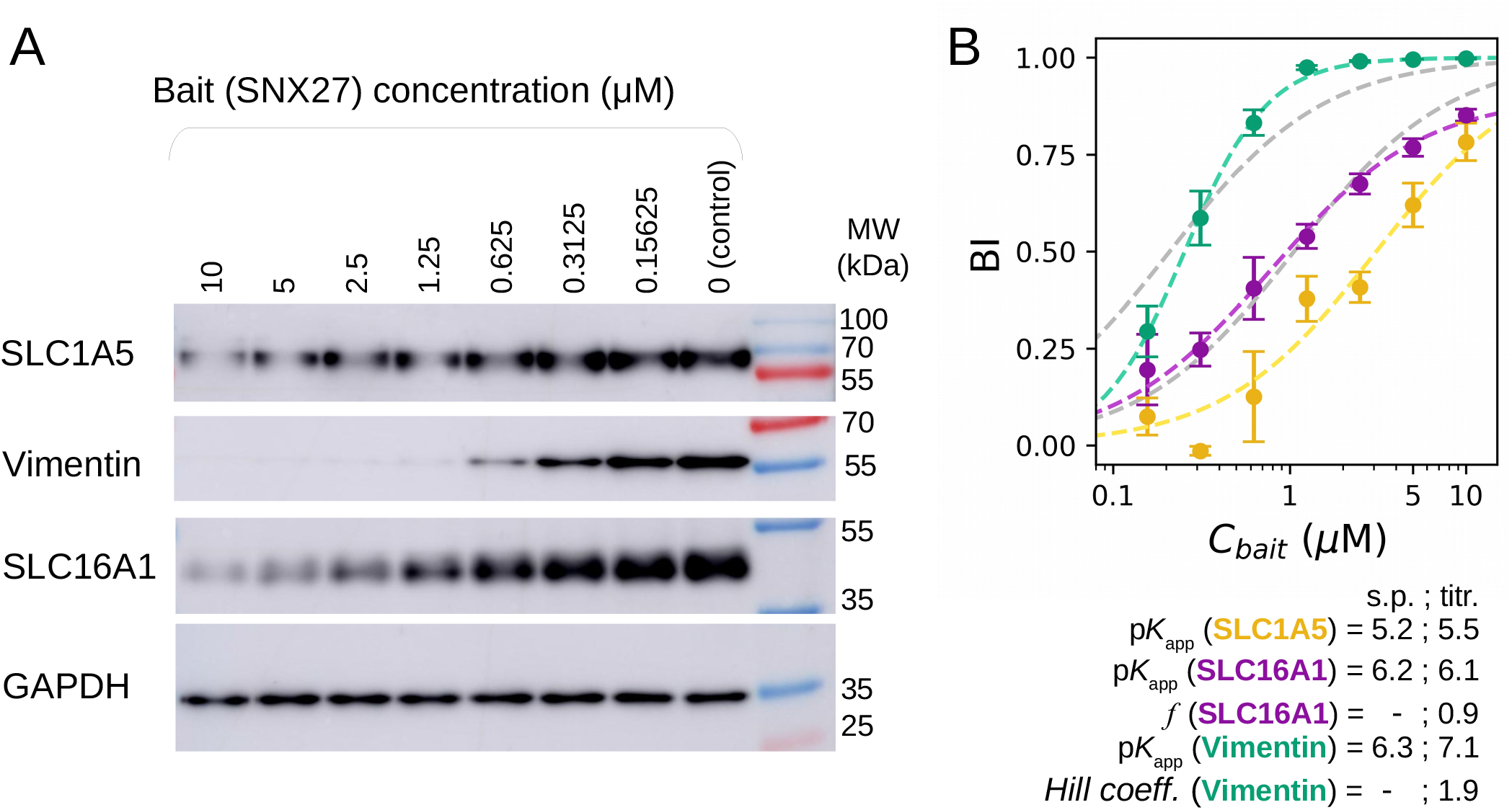
Exploring binding mechanisms of SNX27 interactions with nHU. (A) Results of nHU-WB titration experiments performed with full-length SNX27 bait. (B) Endogenous full-length SLC1A5 (yellow), SLC16A1 (purple) and Vimentin (green) prey depletions were quantified using specific antibodies. BI values of all prey were first fitted with a hyperbolic binding equation. In case of SLC1A5, good fit was achieved with a simple hyperbolic binding equation (yellow dashed line). In case of SLC16A1, an imperfect fit was achieved with a simple hyperbolic binding equation (grey dashed line) and more accurate fit was found when 10% inactive fraction was assumed (purple dashed line, *f* = 0.9, see methods for further details). In case of Vimentin, a low quality fit was achieved with a simple hyperbolic binding equation (grey dashed line) and near perfect fit was obtained using the Hill equation (green dashed line). Equilibrium affinities of single point measurements (s.p.) and parameters determined from the titration experiments (titr.) are indicated below the plots. BI values were determined based on 3 replicates. See Supplementary Figure 3 and Supplementary Table 1 for additional data.

To see if the observed intrinsic binding properties of the above-investigated interactions lead to formation of their complex in cells, we measured co-localization (Figure 6, Supplementary Figure 5). We immuno-stained endogenous interaction partners of SNX27 in U2OS cells expressing HA-tagged full-length or truncated SNX27 lacking its PDZ domain (SNX27_ΔPDZ). Both SNX27 constructs were found to be enriched in endomembrane structures. These SNX27 foci were mostly found in the proximity of Vimentin filaments, however, this was found to be independent of its PDZ-domain, possibly due to the endogenous SNX27 background or due to other interactions (Figure 6E,F). While Vimentin was not enriched in these SNX27 foci, both SLC16A1 and SLC1A5 were (Figure 6A-D). Moreover, both SLC transporters showed statistically significantly stronger co-localization with SNX27 than with SNX27_ΔPDZ. Therefore, these results indicate that while these two transmembrane SLC transporter proteins are cargoes of the SNX27-retromer complex, Vimentin is not and may instead contribute to other activities of the SNX-retromer complex that requires further inspection.

**Figure 6.**
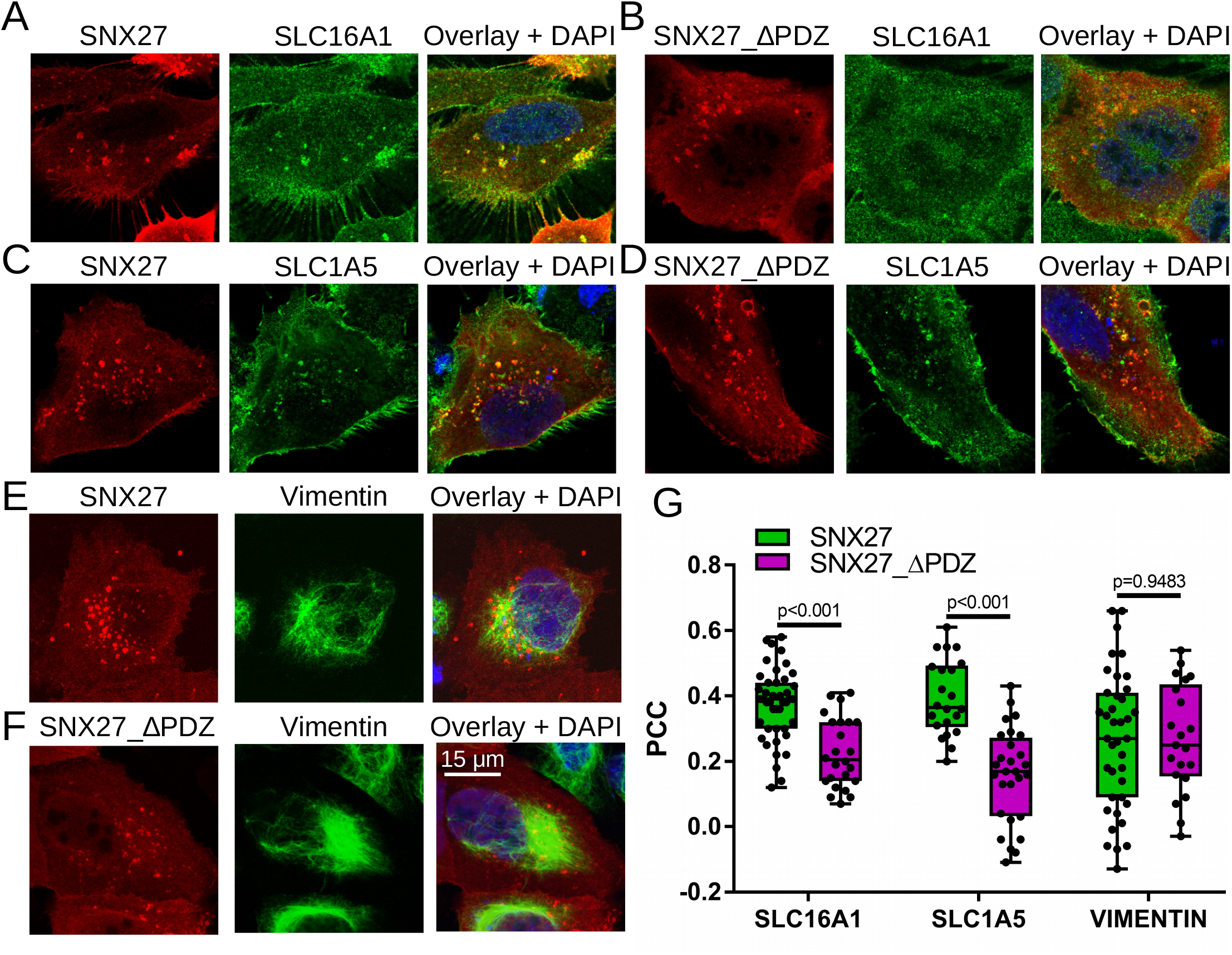
Co-localization of SNX27 and SNX27_ΔPDZ with selected partner proteins identified with nHU. (A-F) Representative co-localization images of U2OS cells expressing HA-tagged SNX27 (A, C, E) or HA-tagged SNX27_ΔPDZ (B, D, F) stained with anti-HA antibody (red) and antibodies against endogenous SLC16A1 (A, B), SLC1A5 (C, D), or Vimentin (E, F) (green), as well as with DAPI (blue). Confocal images are shown for the two transmembrane SLC transporters and a maximum intensity projection is shown for Vimentin. (G) Co-localization was quantified on confocal images for each transfected cell (n > 20) by measuring intensity correlation (PCC). Box plots indicate the median, upper and lower quartiles, and whiskers label the minimal and maximal measured PCC values. Individual data-points representing measurements of single cells are also indicated. P values were calculated using two-sided unpaired T-tests. See Supplementary Figure 5 for additional confocal images.

## Discussion

Although common interactomic assays are efficient for interaction screening, most of them do not measure biophysical properties of interactions. Pulldown-based approaches were used in several ways to gain quantitative insight into affinities of interactions. For example, measured “stoichiometric” ratios of baits and preys have been used in immunoprecipitation experiments as a proxy to discriminate between “strong” and “weak” complexes ^39^. Recently, we have also shown that in parallel pulldown experiments using various baits, the relative enrichment values of endogenous preys correlate with their corresponding fragmentomic affinities ^12^. Pulldown methods were even used to directly estimate affinities of baits using measured enrichment values, yet the consequences of washing steps were not considered ^40 41^. Other types of methods have also been developed to measure affinities directly from cell extracts, but these are mostly low-throughput and require high expertise ^42 43 44 45 46^. Therefore, we still lack a robust method to measure affinities from cell extracts and most HTP affinity measurements are limited to labor-intensive fragmentomic approaches that require expensive reagents and instruments.

Here, we introduced nHU as a versatile tool to measure apparent equilibrium affinities proteome-wide of recombinant or synthetic baits. Although these affinities can be indirect and can be perturbed by protein heterogeneity, nHU experiments can give us insight into an interactomic dimension that was never reached at this scale before: affinities of full-length proteins and even large complexes directly obtained from cell extracts. Compared to qualitative pulldown-based methods, nHU involves less experimental steps and robustly ranks identified targets by their observed affinities. It can be equally used to cost-efficiently screen affinities across the proteome by using single-point measurements, as well as to accurately investigate binding mechanisms using titration experiments. All mature protein analytical technologies can be used in combination with nHU, such as antibody-based approaches like routine western blot or label-free mass spectrometry. In principle, nHU is not limited to studying the interactions of proteins, and both baits and preys can be molecules of different kinds. Overall, nHU experiments can be effortlessly implemented in most laboratories and could greatly advance the exploration of the *quantitative human affinity interactome*.

## Author contribution

All nHU experiments and data analysis were performed by BZ and GG. BZ performed all WB and cell-based assays. MS experiments were performed by BM and LN. GT and GG conceived the idea and supervised the research. BZ and GG wrote the first draft of the paper and all authors reviewed the manuscript.

## Acknowledgements

We are grateful for Kathleen Weimer for proofreeding the first draft of the manuscript. BZ was supported by Fondation pour la Recherche Médicale (FRM, SPF202005011975). GG was supported by the Post-doctorants en France program of the Fondation ARC and by the collaborative post-doc grant of IGBMC. The work was supported by the Ligue contre le cancer (équipe labellisée 2015 to GT), the Agence Nationale de la Recherche (grant UBE3A ANR-18-CE92-0017 to GT), the French Infrastructure for Integrated Structural Biology (FRISBI) ANR-10-INSB-05-01, and Instruct-ERIC.

## Supplementary figure legends

**Supplementary Figure 1.**
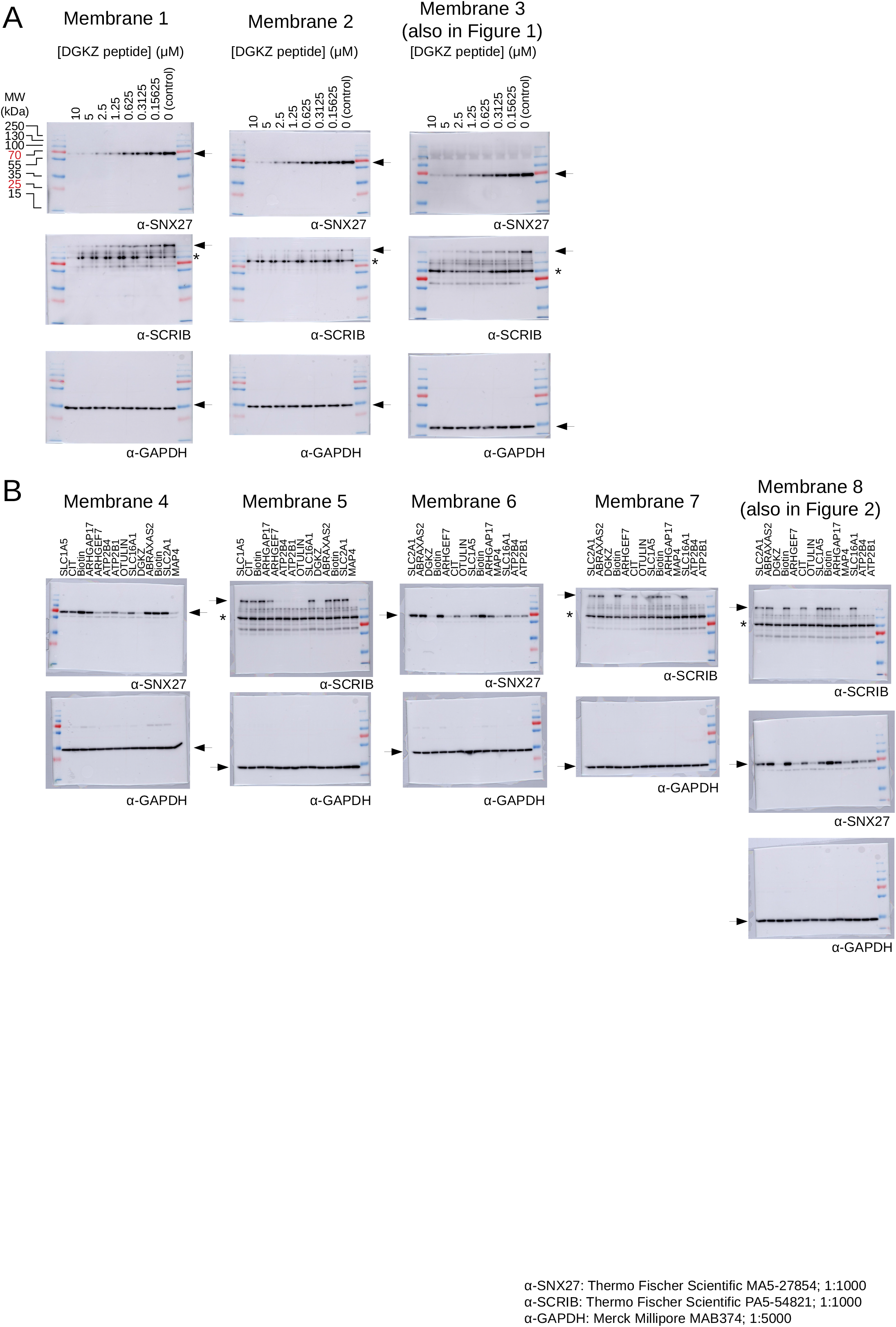
Raw results of nHU-WB experiments using peptide baits. Overlay of luminescent signal (specific signal) with visible image (MW ladder) of all blots used for analysis of nHU-WB experiments using peptide baits. Membranes were H_2_O_2_ treated between SCRIB and SNX27 or SCRIB and GAPDH antibodies. Membranes were mildly stripped between SNX27 and GAPDH antibodies. Asterisk indicates a known aspecific band obtained on western blots labeled with SCRIB antibody. Primary antibodies and their working concentrations are indicated in the bottom right corner. (see Methods for further details)

**Supplementary Figure 2.**
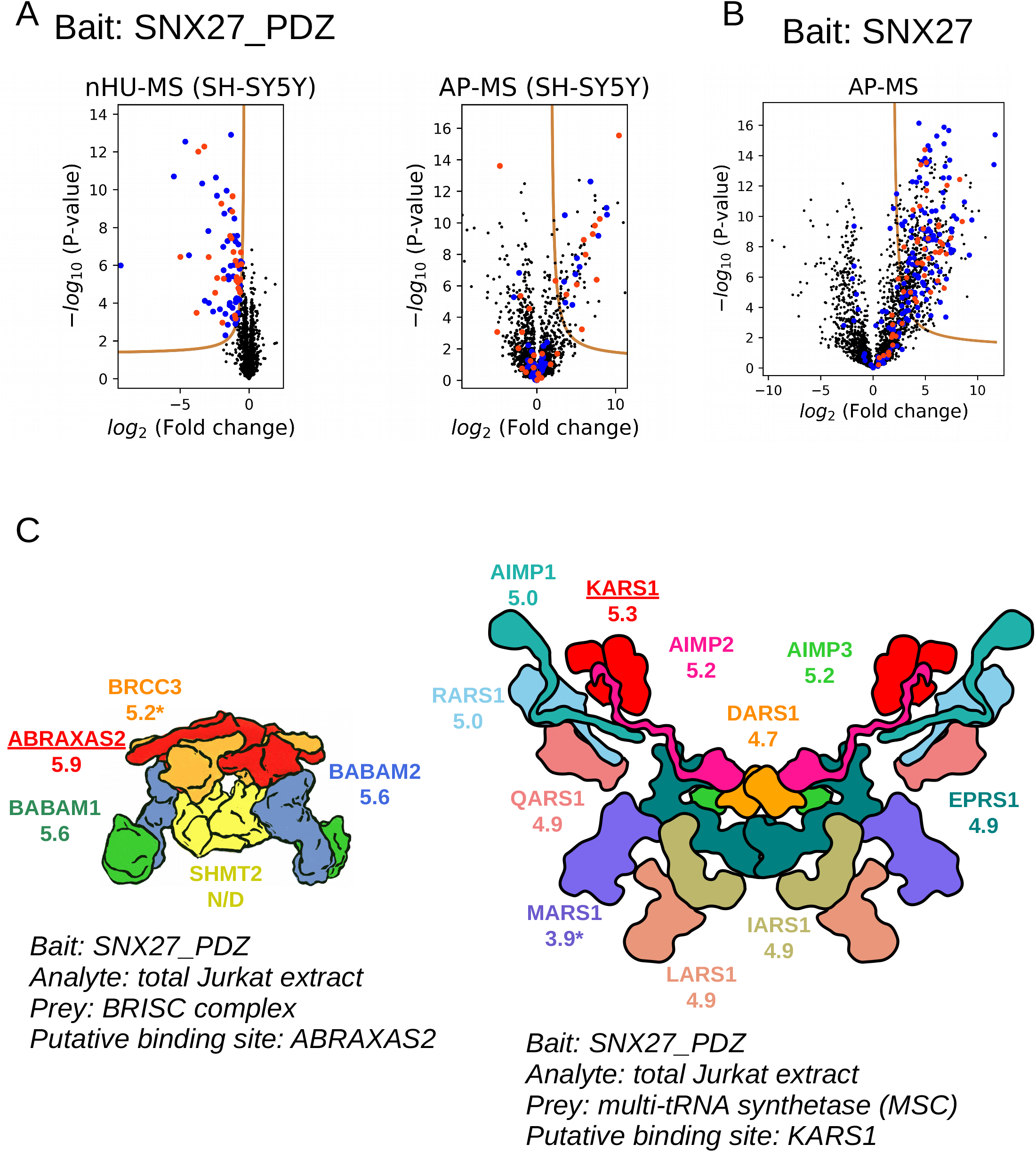
Additional info for nHU-MS experiments. (A) Volcano plot of the nHU-MS (left) and AP-MS (right) experiments performed with recombinant SNX27_PDZ bait and total SH-SY5Y cell extracts (n=6). Converted affinities of the nHU-MS experiments are shown in Figure 3D. (B) Volcano plot of the control pulldown experiment performed on the leftover resin of the nHU experiment showed in Figure 4 (n=6). Preys are colored according to their coloring on Figure 4A. (C) Affinities of two large multi-subunit complexes measured with nHU-MS. The BRISC deubiquitinase complex (left) was found to interact with SNX27 with the exception of the inhibitory subunit SHMT2. Almost all component of the multi-tRNA-synthetase complex (right) was found to interact with SNX27. The subunits with expected direct interactions are underlined on both complexes. For all subunits of the presented complexes, the indicated affinities were obtained with the SNX27_PDZ bait from Jurkat extracts. Asterisk indicates that the affinity was quantified but was found to be not statistically robust. N/D indicates that while the protein amount was quantified, no affinity could be determined from the experiment. See Supplementary Table 1 for additional data.

**Supplementary Figure 3.**
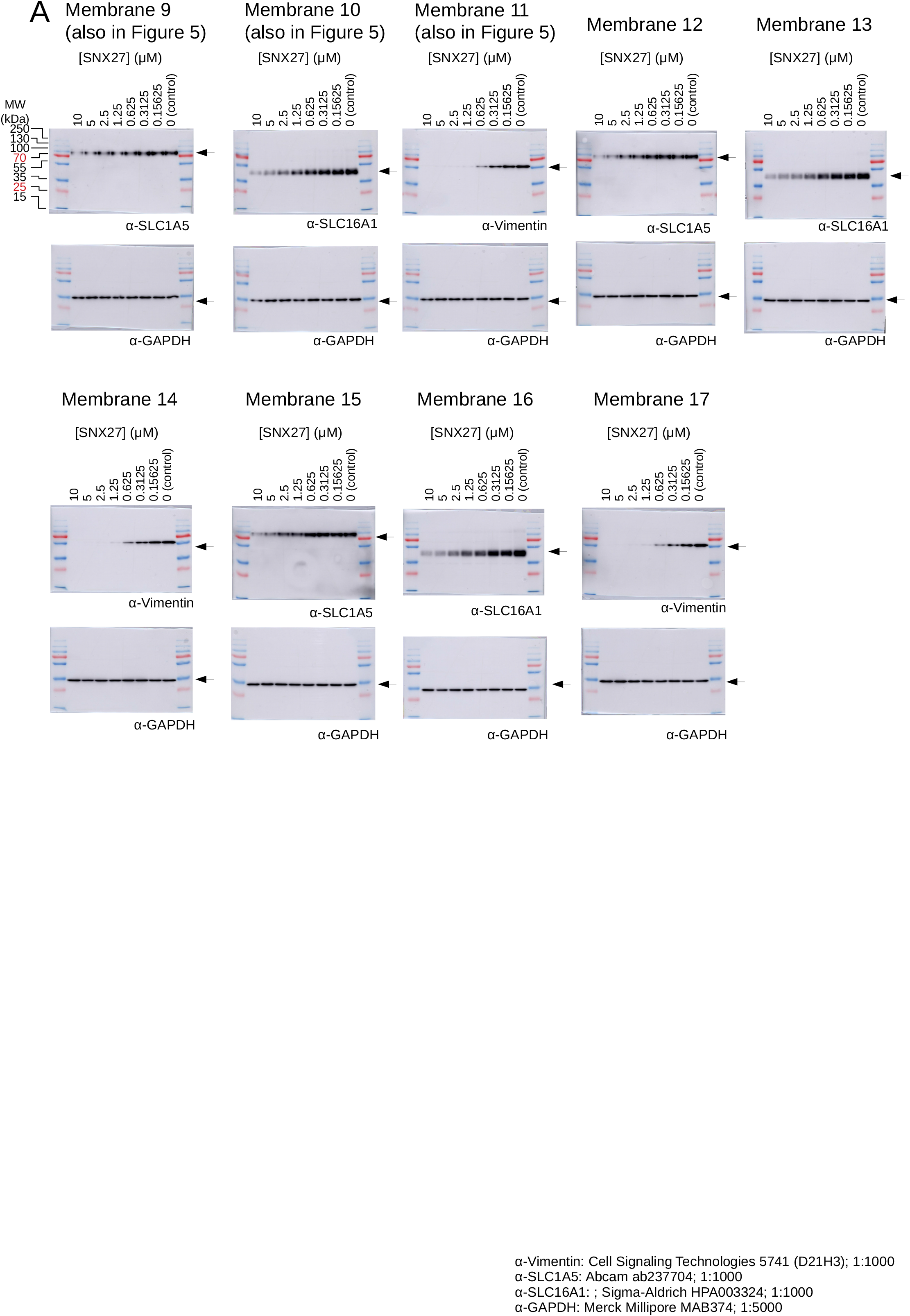
Raw results of nHU-WB experiments using SNX27. Overlay of luminescent signal (specific signal) with visible image (MW ladder) of all blots used for analysis of nHU-WB experiments using SNX27 baits. Membranes were H_2_O_2_ treated before GAPDH blotting. Primary antibodies and their working concentrations are indicated in the bottom right corner.

**Supplementary Figure 4.**
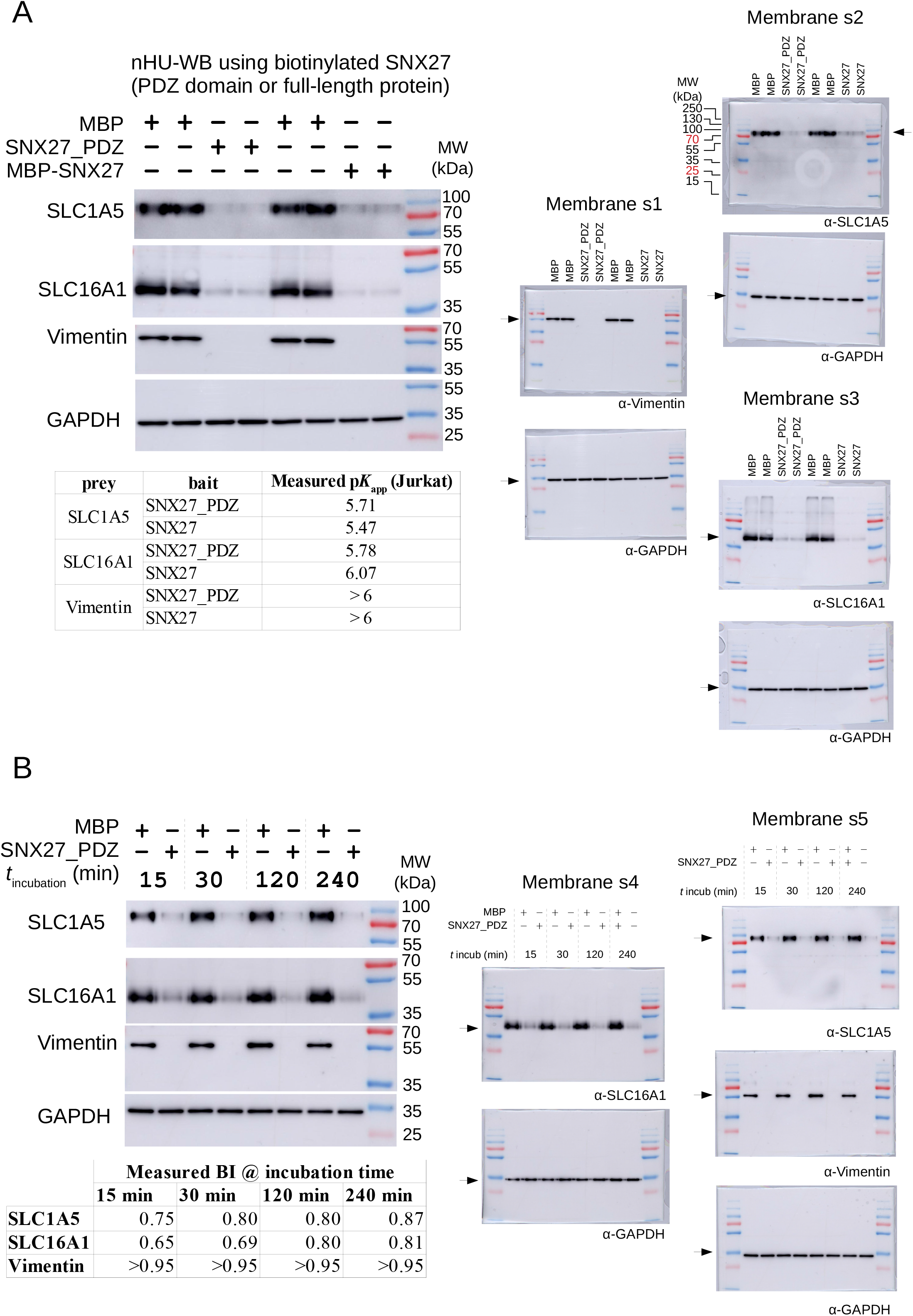
Additional nHU-WB experiments. (A) nHU-WB experiments using SNX27 and SNX27_PDZ baits and total Jurkat extracts. 2 independent experiments are analyzed on the same membrane and the reported affinity was calculated from their averages. (B) nHU-WB experiments performed using SNX27_PDZ bait and Jurkat cell extracts, measured at step-wise increasing incubation times. No detectable change was observed with the different incubation times in the case of Vimentin and only a small change was detected after 30 min incubation in the case of SLC16A1. In all other nHU experiments of our study, 120 min incubation times was applied. Bait concentration was 10 μM in both experimentM in both experiment series. On the right side of both panels, raw results of the western blot experiments are shown similarly to Supplementary Figure 1, or 3. Membranes were H_2_O_2_ treated before GAPDH blotting and mildly stripped between SLC1A5 and vimentin in case of membrane s5.

**Supplementary Figure 5.**
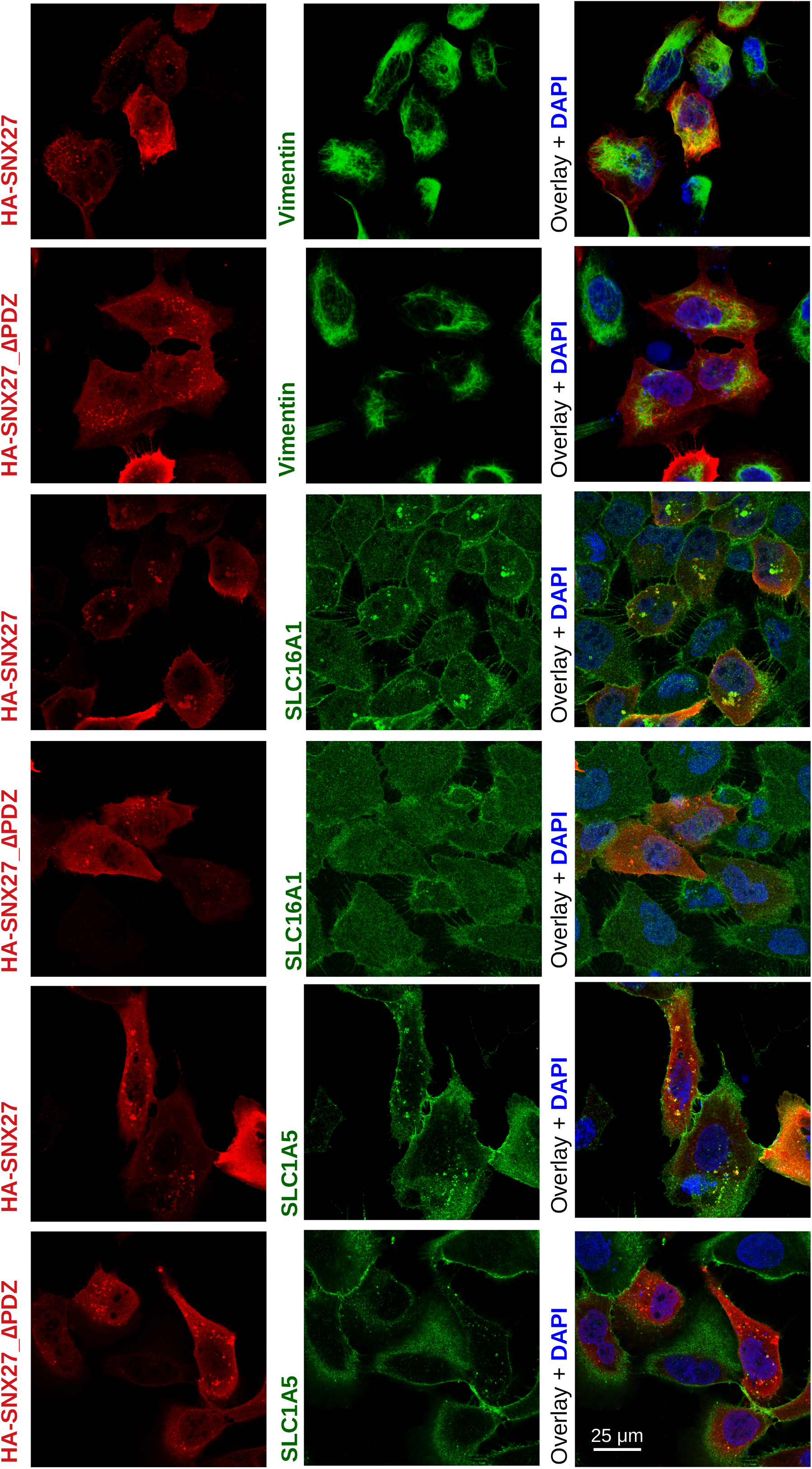
Additional microscopic images for co-localization studies. Representative confocal images of U2OS cells expressing either HA-tagged SNX27 or HA-tagged SNX27_ΔPDZ. Cells were stained with antibodies against HA (red) and endogenous partner protein (Vimentin, SLC16A1, or SLC1A5, all in green), as well as with DAPI (blue).

## Methods

### Peptide and recombinant protein preparation

Biotinylated peptides were chemically synthesized on an ABI 443A synthesizer with standard Fmoc strategy with biotin group attached to the N-terminus via a TTDS (Trioxatridecan-succinamic acid) linker and were purified with HPLC (>95% purity). All C-terminal PBM peptides were 10mer. Predicted peptide masses were confirmed by mass-spectrometry. Peptide concentrations were determined based on their dry weight.

SNX27_PDZ (40-141) was cloned as His_6_-AviTag-MBP-TEV-SNX27_PDZ and full-length SNX27 (1-541) was cloned as His_6_-AviTag-MBP-SNX27-His_6_ construct in pET vectors. The empty His_6_-AviTag-MBP-TEV-vector was used to produce biotinylated MBP for control experiments. Proteins were co-expressed with BirA (PET21a-BirA, Addgene #20857) in *E. coli* BL21(DE3) cells. At IPTG induction (1 mM IPTG at 18 ºC for ON), 50 μM biotin was added to the media. Harvested cells were lysed in a buffer containing 50 mM TRIS pH 7.5, 150-300 mM NaCl, 50 μM biotin, 2 mM BME, cOmplete EDTA-free protease inhibitor cocktail (Roche, Basel, Switzerland), 1% Triton X-100, and trace amount of DNAse, RNAse, and Lysozyme. Lysates were frozen at -20 ºC before further purification steps. Lysates were sonicated and centrifuged for clarification. Expressed proteins were captured on pre-packed Ni-IDA (Protino Ni-IDA Resin, Macherey-Nagel, Duren, Germany) columns, were washed with at least 10 column volume cold wash buffer (50 mM TRIS pH 7.5, 150 mM NaCl, 2 mM BME) before elution with 250 mM imidazole. The Ni-elution was collected directly on a pre-equilibrated amylose column (amylose high flow resin, New England Biolabs, Ipswich, Massachusetts). Amylose column was washed with 5 column volume cold wash buffer before fractionated elution in a buffer containing 25 mM Hepes pH 7.5, 150 mM NaCl, 1 mM TCEP, 10% glycerol, 5 mM maltose, cOmplete EDTA-free protease inhibitor cocktail. The concentration of proteins was determined by their UV absorption at 280 nm before aliquots were snap freeze in LN2 and storage at -80 ºC.

### Cell cultures and extract preparation for nHU experiment

SH-SY5Y cells (ATCC, ref. CRL-2266) were grown in RPMI 1640 (Gibco) medium completed with 10% FCS and 40 μM in both experimentg/mL gentamicin, diluted every 3^rd^/4^th^ day 1/5. U2OS cells (ATCC, ref. HTB-96) were grown in DMEM (Gibco, 1g/l glucose) completed with 10% FCS and 40 μM in both experimentg/mL gentamicin, diluted every 3^rd^/4^th^ day 1/10. Jurkat cells (ECACC) were grown in RPMI (Gibco) medium completed with 10% FCS and 40 μM in both experimentg/mL gentamicin, diluted every 3^rd^/4^th^ day 1/12. All cells were kept at 37°C and 5% CO2.

For preparing semi-native total cell extracts, cells were seeded on T-175 flasks. After they reached confluency, adherent cells were washed with PBS once and collected by scraping with ice-cold lysis buffer (Hepes-KOH pH 7.5 50 mM, NaCl 150 mM, Triton X-100 1%, cOmplete EDTA-free protease inhibitor cocktail 1x, EDTA 2 mM, TCEP 5 mM, glycerol 10%). Jurkat cells were collected by 1000 g x 5 min centrifugation, washed once with PBS, then collected by 1000 g x 5 min centrifugation again and lysed in ice-cold lysis buffer. Lysates were sonicated 4×20 sec with 1 sec long pulses on ice, then incubated rotating at 4°C for 30 min. Lysates were centrifuged at 12,000 rpm 4°C for 20 min and supernatant was kept for further analysis. Total protein concentration was measured by standard Bradford method (Bio-Rad Protein Assay Dye Reagent #5000006) using a BSA calibration curve (MP BIomedicals #160069, diluted in lysis buffer) on a Bio-Rad SmartSpec 3000 spectrophotometer instrument. Lysates were diluted to 2 mg/ml concentration and were snap-freezed in liquid nitrogen and stored at -80°C until measurement.

Note that different lysate preparing protocols can lead to different pools of binding capable proteins and therefore in some cases it may be essential to modify the above described protocol, e.g. by removing EDTA from the lysis buffer in order to measure interactions mediated by metal ions.

### Resin preparation and nHU experiment

For saturating streptavidin resin with biotinylated peptides or biotin, 50 μM in both experimentl streptavidin resin was mixed with biotin or peptide at 40-60 μM in both experimentM concentration in 6-6.5 resin volume for 60 min. For saturating streptavidin resin with biotinylated proteins, 50 μM in both experimentl streptavidin resin was mixed with biotinylated MBP or MBP-PDZ at 40-50 μM in both experimentM concentration in 20 x resin volume for 60 min. After saturation, resins were washed a single time (10 resin volume, holdup buffer: 50 mM Tris pH 7.5, 300 mM NaCl, 1 mM TCEP, .22 filtered), and were depleted with biotin (10 resin volume, 10 min, holdup budder supplemented with 100 μM in both experimentM biotin). Finally, resins were washed 2 times (10 resin volume, holdup buffer).

For single-point nHU experiments carried out at ∼10 μM in both experimentM bait concentration, 50 μM in both experimentl saturated streptavidin resin was mixed with 200 μM in both experimentl 2 mg/ml cell lysate. Titration experiments were carried out by mixing control and bait-saturated resin and keeping the total resin volume constant, e.g. ∼5 μM in both experimentM bait concentration can be achieved by mixing 25 μM in both experimentl bait-saturated streptavidin resin with 25 μM in both experimentl control resin and 200 μM in both experimentl 2 mg/ml cell lysate. Control and bait saturated resins were prepared in larger amounts and a serial dilution was prepared with these pre-saturated resins to achieve different resin ratios.

Unless specified elswhere, the nHU mixture was incubated at 4 ºC for 2 h. After the incubation ended, the resin was separated from the supernatant by a brief centrifugation. Then, half the supernatant was removed by pipetting without any delay to avoid any resin contamination, e.g. 100 μM in both experimentl supernatant is collected if 200 μM in both experimentl cell lysate was used as analyte. Optionally, the supernatant can be centrifuged an additional time to clarify it further removing any possible resin contamination. Note that the separation of the supernatant should be as fast as possible, since any delay can perturb the equilibrium. Alternatively, the resin can be separated from the supernatant using filter plates (e.g. various products of Millipore, Burlington, Massachusetts) or spin columns (e.g. Pierce Spin Cups – cellulose Acetate Filter from Thermo Scientific, Waltham, Massachusetts) to achieve faster separation.

Since exceptionally strong complexes may have extremely slow dissociation rates (*k*_off_) that make it highly difficult to reach binding equilibrium ^24^, we also verified that some interactions indeed reached binding equilibrium using our standard protocol (2 hours of incubation) by probing nHU-WB experiments using various incubation times (Supplementary Figure 4B). We used SNX27_PDZ bait and performed nHU with Jurkat lysates using incubation times of 15, 30, 120, 240 min. Then, nHU supernatants were probed with western blot for SLC1A5, SLC16A1 and Vimentin. For these interaction partners, we did not observe significant change by prolonging the nHU reaction in the monitored timeframe, compared to our standard protocol.

Note that the nHU assay was called as “pure-crude holdup assay from eukaryotic cells” when its first qualitative proof-of-concept was be done ^19^.

### Affinity purification

Leftover beads from nHU-MS experiments were washed 3 times immediately after the separation of the supernatant (10 resin volume in buffer containing: 50 mM TRIS pH 8.5, 150 mM NaCl, 1% TritonX-100, 10x cOmplete EDTA-free protease inhibitor cocktail, 2 mM EDTA, 1 mM TCEP). Then, the beads were washed 2 times (10 resin volume buffer containing: 50 mM TRIS pH 8.5, 150 mM NaCl, 1 mM TCEP). Finally, captured protein was eluted from the resin in two steps and the eluted fractions were pooled. For each elution the beads were incubated for 30 min with 3 resin volume buffer containing: 20 mM TRIS pH8.5, 100 mM NaCl, 500 μM in both experimentM TCEP, 8 M Urea. Between each step, the beads were separated by mild centrifugation and the supernatant was removed by gentle pipetting.

### Western-blot

nHU samples were mixed with 4x Laemmli buffer (120 mM Tris-HCl pH 7, 8% SDS, 100 mM DTT, 32% glycerol, 0.004% bromphenol blue, 1% β-mercaptoethanol) in 3:1 ratio. Equal amounts of samples were loaded on 8% or 10% acrylamide-gels. Transfer was done into PVDF membranes using a Trans-Blot Turbo Transfer System and Trans-Blot Turbo RTA Transfer Kit (BioRad, #1704273). After 1 h of blocking in 5% milk, membranes were incubated overnight 4°C in primary antibody in 5% milk. The following antibodies and dilutions were used: anti-SCRIB (1:1000, Thermo #PA5-54821), anti-SNX27 (1:1000, Thermo #MA5-27854), anti-GAPDH (1:5000, Sigma #MAB374), anti-SLC1A5 (1:1000, abcam #ab237704), anti-SLC16A1 (1:1000, Sigma #HPA003324), anti-VIMENTIN (1:1000, CST #5741). Membranes were washed three times with TBS-Tween and incubated at RT for 1 h in secondary antibody (Jackson ImmunoResearch, Peroxidase conjugated Affinipure goat anti-mouse(H+L) #115-035-146 and goat anti-rabbit(H+L) #111-035-003) in 5% milk (concentration 1:10,000). After washing three times with TBS-Tween, membranes were exposed to chemiluminescent HRP substrate (Immobilon, #WBKLS0100) and revealed in docking system (Amersham Imager 600, GE). Densitometry was carried out on raw Tif images by using Fiji ImageJ 1.53c. Between different primary antibody labeling, the membranes were either exposed to 15% H_2_O_2_ to remove secondary signal (in case of different species) or stripped with mild stripping buffer (15g/L glycine, 1g/L SDS, 1% Tween 20, pH 2.2) to remove primary signal (in case of same species).

### Sample digestion for mass spectrometry

The nHU samples were precipitated with TCA 20% overnight at 4°C and centrifuged at 14,000 rpm for 10 min at 4°C. The protein pellets were washed twice with 1 mL cold acetone and air dried. The protein extracts were solubilized in urea 8 M, reduced with 5 mM TCEP for 30 min and alkylated with 10 mM iodoacetamide for 30 min in the dark. Double digestion was performed at 37°C with 500 ng endoproteinase Lys-C (Wako, Richmond, USA) for 4 h, followed by 4-fold dilution and an overnight digestion with 500 ng trypsin (Promega, Charbonnieres les Bains, France). Peptide mixtures were then desalted on C18 spin-column and dried on Speed-Vacuum.

### LC-MS/MS Analysis

Samples were analyzed using an Ultimate 3000 nano-RSLC (Thermo Scientific, San Jose California) coupled in line with a LTQ-Orbitrap ELITE mass spectrometer via a nano-electrospray ionization source (Thermo Scientific, San Jose California). Peptide mixtures were injected in 0.1% TFA on a C18 Acclaim PepMap100 trap-column (75 μm ID x 2 cm, 3 μm, 100Å, Thermo Fisher Scientific) for 3 min at 5 μL/min with 2% ACN, 0.1% FA in H_2_O and then separated on a C18 Acclaim PepMap100 nano-column (75 μm ID x 50 cm, 2.6 μm, 150 Å, Thermo Fisher Scientific) at 220 nl/min and 38°C with a 90 min linear gradient from 5% to 30% buffer B (A: 0.1% FA in H_2_O / B: 99% ACN, 0.1% FA in H_2_O), regeneration at 5% B. The mass spectrometer was operated in positive ionization mode, in data-dependent mode with survey scans from m/z 350-1,500 acquired in the Orbitrap at a resolution of 120,000 at m/z 400. The 20 most intense peaks from survey scans were selected for further fragmentation in the Linear Ion Trap with an isolation window of 2.0 Da and were fragmented by CID with normalized collision energy of 35%. (TOP20CID method) Unassigned and single charged states were excluded from fragmentation. The Ion Target Value for the survey scans (in the Orbitrap) and the MS2 mode (in the Linear Ion Trap) were set to 1E6 and 5E3 respectively and the maximum injection time was set to 100 ms for both scan modes. Dynamic exclusion was set to 20 s after one repeat count with mass width at ± 10 ppm.

### Mass spectrometry data analysis

Proteins were identified by database searching using SequestHT (Thermo Fisher Scientific) with Proteome Discoverer 2.4 software (PD2.4, Thermo Fisher Scientific) on human FASTA database downloaded from SwissProt (reviewed, release 2021_06_03, 20380 entries, https://www.uniprot.org/). Precursor and fragment mass tolerances were set at 7 ppm and 0.6 Da respectively, and up to 2 missed cleavages were allowed. Oxidation (M, +15.995 Da) was set as variable modification, and Carbamidomethylation (C, + 57.021 Da) as fixed modification. Peptides and proteins were filtered with a false discovery rate (FDR) at 1%. Label-free quantification was based on the extracted ion chromatography intensity of the peptides. All samples were measured in technical triplicates. The measured extracted ion chromatogram (XIC) intensities were normalized based on median intensities of the entire dataset to correct minor loading differences. For statistical tests and enrichment calculations, not detectable intensity values were treated with an imputation method, where the missing values were replaced by random values similar to the 10% of the lowest intensity values present in the entire dataset. Unpaired two tailed T-test, assuming equal variance, were performed on obtained log_2_ XIC intensities. All raw LC-MS/MS data have been deposited to the ProteomeXchange via the PRIDE database with identifier PXD034790.

### Statistics

Since the goal of nHU experiments is to measure affinities instead of identifying interactions partners of high confidence, technical repeats were preferred to measure the occurring changes in protein concentrations more precisely over measuring imprecise affinities from multiple independent nHU experiments. In the case of single point nHU-MS experiments, 2 independent experiments were performed and MS experiments were performed with 3-3 technical replicates (n = 6) and mean intensities were used for fold change calculations. In the case of nHU-WB experiments, western blots were repeated 3 times to minimize the error of densitometric quantification and mean values of determined BI values were used for affinity calculations. Since the same cell extract of known concentration was used for nHU-WB experiments, all samples were handled identically and equal volumes of extracts were loaded on the acrylamide-gels, GAPDH was only used to verify western blot loading qualitatively in nHU-WB experiments. Similar affinities were obtained when densitometric GAPDH levels were used for normalization, however the results showed higher standard deviations, likely because of the inclusion of an additional source of variability.

Measured BI values of nHU-WB experiments were fitted using the hyperbolic binding equation

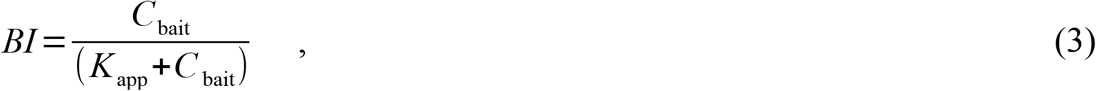

where *C*_bait_ is the immobilized bait concentration and *K*_app_ is the apparent dissociation constant. (Note that eq. 2 is just a rearranged version of eq. 3.) In the case of the nHU-WB experiment between full-length SNX27 and SLC16A1, a BI offset was observed with the hyperbolic fit and a partial activity was assumed using the function:

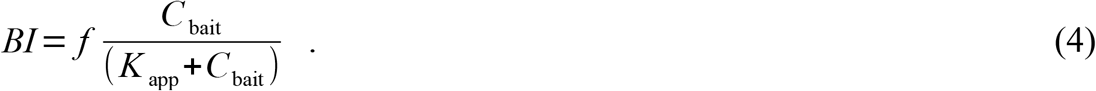

Here, the *f* factor was found to be 0.9, indicating that only 90% of the quantified SLC16A1 protein pool shows binding activity. In cases where the hyperbolic equation could not reach a reasonable solution due to cooperative effects, the Hill equation was used for fitting

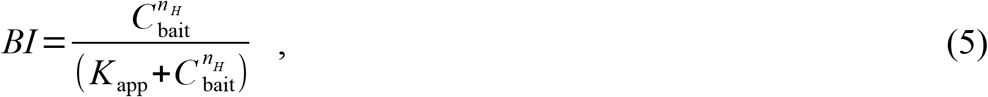

where *n*_H_ is the Hill coefficient (n > 1 for positive and n < 1 for negative cooperative interactions). Fitting was performed in QtiPlot using standard procedures (Scaled Levenberg-Marquardt algorithm with 1000 iterations) and figures were generated with custom Python scripts using Matplotlib.

Affinities for each detected proteins were calculated in nHU-MS experiments where the experimental BI was a positive value using equation 2, however only those were considered for subsequent analysis where statistically robust was observed. For each detected protein in nHU-MS experiments, a P value was calculated based on the measured intensities of samples (n=6) and controls (n=6) using a two-tailed unpaired Student’s T-test. For each nHU-MS and AP-MS experiment series, a hyperbolic binding threshold was calculated taking into account both measured intensities and the general distribution of the entire dataset, similarly as described in other works ^4^. This threshold was calculated for AP-MS as follows:

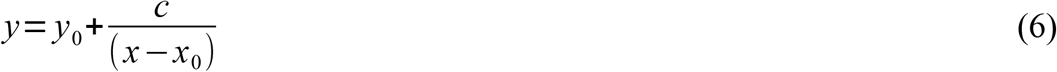

and for nHU-MS experiments as:

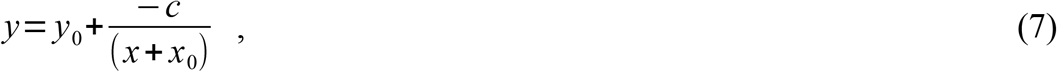

where *y* is the P value threshold at fold change of *x, c* is a curvature parameter empirically fixed at 1 for nHU-MS and 4 for AP-MS experiments, *y*_0_ is the minimal P value threshold and *x*_0_ is the minimal fold change value for any given dataset. The minimal P value was defined at 1.3 -log_10_(P) and thus there is a >95% probability that all identified interaction partners are true interaction partners. The minimal fold-change value cutoff was set at 1 σ and was determined by measuring the width of the normal distribution of all measured fold changes in a given experiment. Note that this threshold can be only interpreted for interaction partners with fold change values of -*x > x*_0_ in case of nHU-MS experiments or *x > x*_0_ in case of AP-MS experiments.

Although standard deviations of measured BI values can be calculated from replicates, it is challenging to propagate these errors to affinities. While BI measurements define the overall precision of any holdup assay, the affinity accuracy mainly depends on the bait concentration. At the moment, this parameter is only estimated based on previous observations ^12^. In future generations of the nHU, more direct methods are needed to measure the error of this estimation in the future that can be used to calculate affinities at high accuracy. For these reasons, we do not specify propagated errors for the calculated affinities since they would be misleading because absolute affinities could change with a more accurate bait concentration. However, affinity ranking of different prey measured from the same nHU experiment will be unaffected by such transformation. For example, based on our experiments we can conclude that the true affinity of SNX27 with Vimentin is stronger than with SLC16A1 and the affinity with SLC16A1 is stronger than with SLC1A5, but it is possible that these affinities can diverge systematically compared to the reported affinities.

### Calculating amounts of complex formations

Determined affinities can be combined with total protein concentrations in order estimate amounts of binary complexes. These coarse predictions can be performed for any cellular proteomes, even at subcellular resolutions. We performed such calculations to estimate amounts of SNX-retromer-WASH complexes bound to SNX27 using estimated protein concentrations previously measured for HEK293T cells ^35^. In principle, such absolute proteomic datasets could be also estimated from the control nHU experiments directly using straightforward tools such as the proteomic ruler, however one has to assume that the prepared cellular extract is representative for the entire cellular proteome of the monitored cell. In these calculations, one can not use the hyperbolic binding equations (eq. 2 or eq. 3) since the concentrations of binding partners are comparable and the partial binding occupation of SNX27 can’t be neglected. Instead, we used the quadratic binding equation (or similar) to make predictions:

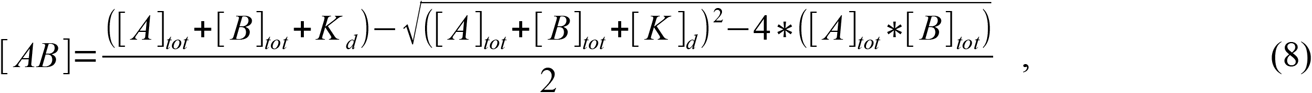

where [AB] is the concentration of the complex under equilibrium, [A]_tot_ and [B]_tot_ are the total concentrations of the binding partners (e.g. quantified with absolute proteomics) and K_d_ is the steady-state dissociation constant. Calculated amounts of complexes were also converted into % of SNX27 bound by comparing the amounts of complexes with the total SNX27 concentration (280 nM). Note that although such calculations can be performed for PDZ-mediated interactions, results will be flawed due to their mutually exclusive nature. Future rule-based network-level calculations should consider affinities and concentrations of all PDZ and PBM proteins, as well as their binding mechanisms, to estimate amounts of all possible complexes in the network.

### Immunostaining

For transient transfection, the full-length SNX27 (1-541) or SNX27_ΔPDZ (140-541) constructs were cloned in mammalian pCI vector containing N-terminal HA tag for immuno-labelling. For detection of protein localization, 0.25×10^5^ U2OS cells per well were seeded onto a cover slip-containing 24-well plate. The next day, cells were transfected with HA-tagged constructs using JetPRIME reagent (Polyplus) and 24h after transfection were washed once and fixed for 15 min with 4% PFA solution, permeabilized for 10 min with 0.3% Triton-X in PBS at room temperature and blocked for 1 h in 5% BSA and 0.3% Triton-X in PBS at room temperature. Staining of HA, SLC1A5, SLC16A1, Vimentin was performed overnight at 4°C using anti-HA (1:750, BioLegend #901502), anti-SLC1A5 (1:200, abcam #ab237704), anti-SLC16A1 (1:200, Sigma #HPA003324), anti-Vimentin (1:500, CST #5741), respectively. After three washings with PBS, secondary antibodies were used for 1 hour at room temperature in 5% BSA and 0.3% Triton-X in PBS: Alexa Fluor 594 conjugated anti-mouse (1:1000; Invitrogen #A-11032) and Alexa Fluor 488 conjugated anti-rabbit (1:1000; Invitrogen #A-11034). Cover glasses were mounted to microscopy slides by Vectashield mounting medium with DAPI (Vector laboratories, Burlingame, California). Images were taken using a Leica SP5 confocal microscope (Leica Camera AG, Wetzlar, Germany) with a HCX PL APO 63×/1.40-0.60 OIL objective using excitation at 405 nm (diode), 488 nm (Argon laser), and 594 nm (HeNe laser) and emission at 415-480, 510-560, 610-695 nm, for DAPI, Alexa488, and Alexa594, respectively. Images were processed by the Fiji ImageJ software. In every images, transfected cells based on HA signal were selected as ROIs manually and Coloc 2 plugin was used to determine single-cell PCC values. Statistics and box plots were done using GraphPad Prism 7 software.

